# Analog of Kynurenic Acid Decreases Tau Pathology by Modulating Astrogliosis in Rat Model for Tauopathy

**DOI:** 10.1101/2022.04.19.488739

**Authors:** Petra Majerova, Dominika Olesova, Greta Golisova, Martina Buralova, Alena Michalicova, Jozef Vegh, Juraj Piestansky, Mangesh Bhide, Jozef Hanes, Andrej Kovac

**Affiliations:** Institute of Neuroimmunology, Slovak Academy of Sciences, Dubravska cesta 9, 845 10 Bratislava, Slovakia; Faculty of Natural Sciences, Department of Biochemistry, Comenius University in Bratislava, Mlynska dolina, Ilkovicova 6, 842 15 Bratislava, Slovakia; Department of Pharmaceutical Analysis and Nuclear Pharmacy, Faculty of Pharmacy, Comenius University in Bratislava, Odbojarov 10, SK-832 32 Bratislava, Slovakia; Laboratory of Biomedical Microbiology and Immunology, The University of Veterinary Medicine and Pharmacy in Kosice, Kosice, Slovakia

**Keywords:** Kynurenic acid, tau, Alzheimer’s disease, inflammation, astrocytes

## Abstract

**Background and purpose:** Kynurenines have immunomodulatory and neuroactive properties and can influence the central nervous system. Previous studies showed the involvement of the kynurenines in the pathogenesis and progression of neurodegenerative disease. In neurodegenerative disorders, including tauopathies, the tryptophan metabolism is shifted toward neurotoxic agents and the reduction of neuroprotectant products. Astrocyte-derived kynurenic acid serves as a neuroprotectant. However, systemic administration of kynurenic acid is not effective because of low permeability across the blood-brain barrier (BBB).

**Experimental Approach:** We used a kynurenic acid analog with similar biological activity but higher brain permeability to overcome BBB limitations. In the present study, we used amide derivate of kynurenic acid N-(2-N, N-dimethylaminoethyl)-4-oxo-1H-quinoline-2-carboxamid (KYNA-1). We administered KYNA-1 for three months to tau transgenic rats SHR-24 and analyzed the effect on tau pathology and activation of glial cells. Primary glial cell cultures were applied to identify the mechanism of the KYNA-1 effect.

**Key results:** KYNA-1 was not toxic to rats after chronic three-month administration. When chronically administered, KYNA-1 reduced hyperphosphorylation of insoluble tau in the brain of transgenic rats. Noteworthily, the plasma total tau was also reduced. We determined that the effect of KYNA-1 on tau pathology was induced through the modulation of glial activation. KYNA-1 inhibited LPS induced activation of astrocytes and induced transformation of microglia to M2 phenotype.

**Conclusion and Implications:** We identified that the administration of KYNA-1 reduced tau hyperphosphorylation and neuroinflammation. KYNA-1 may serve as a promising treatment for tauopathies.

**What is already known?:**

- Studies showed tryptophan-kynurenine pathway changes in neurodegenerative disorders including tauopathies
- Kynurenines exert immunomodulatory and neuroactive properties and have influence on the central nervous system

**What does this study add?:**

- Chronic administration of synthetic analog of kynurenic acid (KYNA-1) reduces tau phosphorylation and astrogliosis in a transgenic rat model for tauopathies
- The analog reversed LPS-induced inflammatory changes in glial cell cultures

**What is the clinical significance?:**

- Administration of KYNA-1 analog shifted the tryptophan metabolism in the neuroprotectant direction
- Neuroprotective analogs KYNA-1 can serve as a new and effective potential therapeutic approach for tauopathies

## 1 INTRODUCTION

Tauopathies are a heterogeneous group of neurodegenerative disorders, pathologically characterized by neuronal and/or glial intracellular inclusions of the microtubule-binding protein, tau. The pathogenesis of tauopathies has been associated with systemic inflammation and disruption of the serotonergic signaling system. A common link between neuroinflammation and disruption of the serotonergic signaling system is the catabolism of the essential amino acid-L-tryptophan [1]. The majority of tryptophan is metabolized via the enzymes tryptophan 2,3-dioxygenase and indoleamine 2,3-dioxygenase in the kynurenine pathway (KP) [2]. The KP generates a series of catabolites collectively known as kynurenines. In the KP, tryptophan is metabolized to kynurenine, which is transformed by kynurenine aminotransferase to kynurenic acid (KA) and by kynurenine hydroxylase to 3-hydroxykynurenine, which is further metabolized to quinolinic acid, the precursor of nicotinamide adenine dinucleotide (NAD^+^) [3].

Kynurenines exert immunomodulatory and neuroactive properties and can influence the functioning of central nervous system (CNS) [4]. Several studies have led to the hypothesis that microglial-derived 3-hydroxykynurenine and quinolinic acid harbor neurotoxic functions while astrocyte-derived KA has neuroprotective functions. In neurodegenerative conditions, the tryptophane metabolism is directed toward neurotoxic agents and there is a significant reduction of neuroprotectant products. Therefore, the shift of the tryptophan metabolism in the neuroprotectant direction may serve as a new and effective therapeutic approach [5].

In the brain, KA selectively acts as an antagonist of ionotropic glutamate N-methyl-D-aspartate (NMDA) receptors by blocking the co-agonist site for glycine, as well as a non-competitive inhibitor of α7-nicotinic receptors for acetylcholine. KA controls amounts of neurotransmitters, such as glutamate, acetylcholine, dopamine, and γ-aminobutyric acid (GABA) and protects neuronal cells from excitotoxicity induced by the overactivation of NMDA receptors [6]. It could modulate the neurotoxic effects of quinolinic acid. Furthermore, KA also plays a role in pathological states, including inflammatory, vascular, and antioxidant processes. Anti-inflammatory functions of KA are of particular interest. Its anti-inflammatory effects were observed in cell models and further confirmed by *in vivo*studies in mice. KA treatment inhibited the lipopolysaccharide (LPS) induced increase of tumor necrosis factor-α (TNF-α) and nitric oxide (NO) in mice serum and also significantly reduced LPS-induced death in animals [7].

Dysregulated metabolism of KA has been found in several neurological and neurodegenerative diseases including Huntington’s disease, Parkinson’s and Alzheimer’s disease (AD), amyotrophic lateral sclerosis, multiple sclerosis, and migraine [8-11]. Several studies indicate the significant involvement of the KP in the pathogenesis and progression of AD. Besides that, the kynurenine/tryptophan ratio has been proposed as a biomarker for detecting systemic inflammation in AD and correlated with the cognitive performance of the patients [12]. The serum and red blood cell KA levels were decreased in AD patients. On the other hand, significantly increased levels of KA were found selectively in the caudate nucleus and the putamen. Another study demonstrated lower KA concentration in the lumbar cerebrospinal fluid (CSF) in AD patients [13].

The manipulation of the KP balance through the levels of KA has become an attractive target for the development of pharmacological agents with the potential to treat neurodegenerative and neurological disorders [14]. However, KA is not able effectively cross the blood-brain barrier (BBB) and thus cannot be diretly used as a neuroprotective agent [15]. Novel strategies are needed to deliver KA into the brain, including vector-mediated drug delivery systems or synthetic approaches such as the chemical modifications of KA. Several new analogs of KA, prodrugs, or derivates with a similar biological effect but improved ability to cross BBB were prepared recently [16, 17]. The prodrug 4-chlorokynurenine (4-Cl-KYN) can be readily transported through the BBB into the brain where produces high levels of 7-chlorokynurenine (7-Cl-KYN), a highly selective antagonist of the NMDA receptor. The chronic administration of 4-Cl-KYN reduced the kainate-induced seizure activity and reduced the neurotoxic effect of quinolinic acid in the rat hippocampus and striatum [18, 19]. Several analogs of KA are in clinical trials for stroke, chronic pain and multiple sclerosis [20].

The new analog N-(2-N,N-dimethylaminoethyl)-4-oxo-1H-quinoline-2-carboxamide hydrochloride (KYNA-1) has proved to be neuroprotective in several models and exhibits feature similar to those of KA. In the micromolar range, its administration decreased the amplitude of the field excitatory postsynaptic potential potentiation in the hippocampus [21-23]. However, preclinical and subsequent clinical investigations of KYNA-1 are needed to evaluate its usefulness in neurodegenerative diseases such as tauopathies.

In this study, we for the first time tested the effect of KYNA-1 on the progression of neurofibrillary pathology in tauopathies using a transgenic animal model SHR-24. We describe the *in vitro*BBB permeability of KYNA-1, plasma, and brain pharmacokinetics after single bolus administration, impact on levels of KP metabolites, and activation of astrocytes and microglia. We report that chronic administration of KYNA-1 reduces the tau pathology in the brain through the modulation of astrogliosis.

## 2 METHODS

### 2.1 Chemicals and materials

Analytical standards anthranilic acid (AA), kynurenic acid (KA), xanthurenic acid (XA), and L-kynurenine (L-KYN) were purchased from Sigma-Aldrich (St. Louis, MO, USA). Internal standards AA (3,4,5,6-D4) and L-KYN (D4) were purchased from Buchem B.V. (Apeldoorn, Netherlands), XA (D4) from Santa Cruz Biotechnology, Inc. (Texas, USA), KA (3,5,6,7,8-D5) from CDN isotopes (Quebec, Canada). KYNA-1 and its internal standard were synthesized by SilharCo s.r.o. (Kamenicany, Slovakia), Dimethylsulfoxide (DMSO), trichloroacetic acid (TCA), formic acid (FA), and LC-MS grade acetonitrile were purchased from Sigma-Aldrich (St. Louis, MO, USA). All standards were of analytical grade. Water was purified using a Millipore purification system (Bedford, MA, USA).

### 2.2 Animals

All animals used in this study were from the in-house breeding colony (SPF-like, monitored according to the Federation of European Laboratory Animal Science Associations). For pharmacokinetic study and isolation of primary rat endothelial cells, we used Sprague-Dawley (SD) rats (6 months old). For chronic administration of KYNA-1 we used 6 months old SHR-24 trangenic rat model for tauopathy with progressive neurofibrillary pathology in the brainstem and frontal cortex [24]. For isolation of primary rat astrocytes, we used 1-2 days old SD rats. All experiments on animals were carried out according to the institutional animal care guidelines conforming to international standards (Directive 2010/63/EU) and were approved by the State Veterinary and Food Committee of Slovak Republic (Ro-933/19-221/3A). Each animal was monitored daily (food consumption, water consumption, and overall health status). Humane endpoints based on body weight loss and body condition scoring were in place. For terminal experiments, animals were euthanized with CO_2_. No unexplained mortality occurred in these studies.

### 2.3 Isolation and cultivation of primary rat glial culture

Rat mixed glial culture was prepared from cerebral cortices of 1–2 days old SD rats (n = 8/isolation). The animals were euthanized by CO_2,_and the cerebral cortices were dissected, stripped of the meninges, and mechanically dissociated by repeated pipetting followed by passage through a 20 μm nylon mesh (BD Falcon, Franklin Lakes, USA). Cells were plated on 6-well plates pre-coated with poly-L-lysine (10 μg ml^-1^, Sigma-Aldrich, St. Louis, MO, USA) and cultivated in DMEM medium (PAA Laboratories GmbH, Germany) containing 10% fetal calf serum (FCS, Thermo Fisher Scientific, Waltham, Massachusetts, USA) and 2 mM L-glutamine (Life Technologies Invitrogen, Carlsbad, CA) at 37°C, 5% CO_2_in a water-saturated atmosphere. For the experiments, we used 3 weeks old glial cells. Astrocyte cultures were prepared by mild trypsinization.

### 2.4 Development of multi-component cell model system

The multi-component model was composed of primary rat astrocytes and human neuroblastoma cell line SH-SY5Y with inducible expression of truncated tau protein (aa 151-391/4R) [25]. Firstly, the expression of truncated tau protein was induced by the cultivation of cells in a medium without doxycycline (Sigma-Aldrich, St. Louis, MO, USA) for 3 days before cell seeding into co-culture. Secondly, SH-SY5Y cells were replated into primary astrocytes in 6-well plates in density 1.10^3^cells/cm^2^and co-cultivated for further 5 days in DMEM medium (PAA Laboratories GmbH, Germany) containing 1% FCS (Invitrogen, Carlsbad, CA), 2 mM L-glutamine (PAA Laboratories GmbH, Germany) and N2-supplement (Life Technologies Invitrogen, Carlsbad, CA) at 37°C and 5% CO_2_. The medium was changed twice a week.

### 2.5 Isolation and cultivation of primary rat brain endothelial cells (RBECs)

Isolation of primary brain endothelial cells was performed as previously described [26]. Briefly, rats (200–250 grams, 6 months old; n = 4/isolation) were euthanized by CO_2,_and their brains were removed. Under sterile conditions, the brainstem and cerebellum were dissected, and white matter from the midbrain and the choroid plexus were removed. The cortical tissues were cleansed from meninges and were homogenized on ice in DMEM-F12 medium (PAA Laboratories GmbH, Germany) with 0.1% bovine serum albumin (BSA, Sigma-Aldrich, St. Louis, MO, USA). The homogenate was centrifuged at 800 x g for 10 min at 4°C. The supernatant was aspirated and the pellet was resuspended in a pre-warmed digestion mix containing 1 mg ml^-1^collagenase/dispase (Roche Diagnostics, Indianapolis, USA) and 10 μg ml^-1^DNase I (Roche Diagnostics, Indianapolis, USA). The homogenates were incubated with a pre-prepared digestion mix at 37°C for 30 min with gentle shaking. The preparation was centrifuged at 800 x g for 10 min at 4°C, and the pellets were resuspended in 20% BSA in the medium. The tissues were centrifuged at 1500 x g for 15 min at 4°C to obtain pellet containing microvessels with a fraction of myelin and BSA on the top which was centrifuged again. The microvessels were pooled and resuspended in a pre-warmed digestion mix and incubated for 15 min at 37°C. The pellet was centrifuged at 800 x g for 10 min at 4°C and washed with a serum-containing DMEM-F12 culture medium. The microvessels were cultured in DMEM-F12 medium containing 15% plasma-derived serum (PDS, First Link, UK), 2 mM UltraGlutamine (GE Healthcare, UK), BME vitamins (Sigma-Aldrich, St. Louis, MO, USA), heparin (Sigma-Aldrich, St. Louis, MO, USA) and 3 μM puromycin (Sigma-Aldrich, St. Louis, MO, USA).

### 2.6 Development of *in vitro*model of BBB

After 7 days, endothelial cells were plated onto the top of 0.4 μm Transwell inserts (Becton Dickinson, New Jersey, USA) pre-coated with 10 μg/cm^2^collagen type IV (Sigma-Aldrich, St. Louis, MO, USA) and 5 μg/cm^2^fibronectin (Sigma-Aldrich, St. Louis, MO, USA). The cells were cultivated together with mixed glial cells plated on the bottom of 12-well plates for 7 days in EBM-2 medium (Lonza, Basel, Switzerland) containing 15% PDS, 2 mM UltraGlutamine, BME vitamins, and BulletKit SingleQuots (Lonza, Basel, Switzerland). The TEER values of *in vitro*BBB model were determined by Ohm meter “EVOM” (World Precision Instrument, EVOM Sarasota, FL, USA).

### 2.7 Permeability assay

For the permeability experiments, we used only inserts with TEER values higher than 300 Ω.cm^2^. Transwell inserts were washed with Ringer-HEPES buffer (150mM NaCl, 5.2mM KCl, 2.2mM CaCl_2_, 0.2mM MgCl_2_·6H_2_O, 6mM NaHCO_3_, 5mM HEPES, 2.8mM glucose; pH 7.4). KYNA-1 was dissolved in Ringer-HEPES and applied in the upper (apical) compartment in a final concentration of 1mg ml^-1^. All incubations were performed at 37 °C. Samples were taken from the basolateral side after 5, 15, 30, and 60 min. The concentrations of KYNA-1 in abluminal and luminal compartments were measured by the validated LC-MS/MS method. The permeability values of the inserts (PSf, for inserts with a coating only) and the insert plus endothelium (PSt, for inserts with a coating and cells) were taken into consideration by applying the following equation: 1/PSe = 1/PSt − 1/PSf. To obtain the endothelial permeability coefficient (Pe, in cm/s^-1^), the permeability value (PSe) corresponding to the endothelium alone was then divided by the insert’s porous membrane surface area.

### 2.8 Determination of IC_50_

The cell toxicity was determined by measuring the relative amount of adenylate kinase released from cells into the medium (ToxiLightTM Non-Destructive Cytotoxicity BioAssay Kit, LONZA, Swittzerland). Medium from cell culture was mixed with adenylate kinase substrate adenosine diphosphate. Then luciferin and luciferase were added and the resulting bioluminescence was measured using a luminometer Fluoroscan Ascent FL (MTX Lab Systems, Inc.). Cytotoxicity data were fitted to a sigmoidal curve and a four-parameter logistic model was used to calculate IC_50_. The analysis was performed using Prism 8.0 (GraphPad Inc, San Diego, CA, USA).

### 2.9 Cell stimulation and nitrite assay

*In vitro*neuroinflammation was induced by 250ng ml^-1^LPS from E. coli; O26:B6 (Sigma-Aldrich, St. Louis, MO, USA) to primary rat astrocytes for 24 hours. Nitrite, a downstream product of NO, was measured by the Griess reaction in culture supernatants as an indicator of NO production. Briefly, 50µl of cell culture medium was incubated with 100µl of Griess reagent A (1% sulfanilamide (Merck, Darmstadt, Germany) and 5% phosphoric acid (Slavus, Bratislava, Slovakia) in distilled water for 5 min, followed by the addition of 100µl Griess reagent B (0,1% N-(1-naphthyl) ethylenediamine (Honeywell, North Carolina, USA) in distilled water for 5 min. The absorbance was measured at 540 nm using a microplate reader (PowerWave HT; Bio-Tek).

### 2.10 KYNA-1 formulation for administration

The intraperitoneal formulation was a solution prepared by dissolving KYNA-1 in a sterile sodium chloride physiological solution. The solution was vortexed for 3 min and filtered with a 0.22 μm nylon syringe filter (Pall Corporation, NY, USA). KYNA-1 was administrated intraperitoneally (i.p.) in a concentration of 200 mg kg^-1^of body mass.

### 2.11 In vivo preclinical pharmacokinetic studies

The pharmacokinetic studies of KYNA-1 were performed in SD rats (bodyweight 300 ± 50 g). The SD rats starved 12 h before dosing. For the pharmacokinetic analyses, the animals were randomly divided into seven groups. For each time point, we used eight rats per group. A single dose of 200 mg kg^-1^KYNA-1 was intraperitoneally administered to the animals. The time points for the dosing groups were at pre-dose and post-dose (0.5, 1, 2, 6, 12, 24, 48 h) of the KYNA-1. After dosing, brain tissue was removed, and blood samples were collected. The concentration of KYNA-1 was measured using LC-MS/MS. Plasma pharmacokinetic parameters of KYNA-1 post i.p. administration were calculated using the PKSolver 2.0 software [27]. The brain tissue concentrations were corrected for residual blood [28]. The brain/serum ratios were plotted against exposure time using Prism 6.0 (GraphPad Inc, San Diego, CA, USA). The unidirectional influx rate (*K*_i_in μl g^−1^min^−1^) and *V*_i_, the vascular space, and initial luminal binding at *t*= 0 were calculated from the curve.

### 2.12 Rat plasma and brain tissue collection

Plasma was obtained from all rats using a cardiac puncture protocol. Briefly, the animals were anesthetized with tiletamine-zolazepam/xylazine anesthesia, and cardiac puncture was rapidly performed; the blood samples were then collected into heparinized tubes. Plasma was harvested by centrifuging the blood samples at 8 000 rpm for 15 min and stored at -80°C until analysis. The brain tissue was removed and dissected to the brainstem and cortex. The tissue samples were frozen in liquid nitrogen and stored at -80°C until analyzed.

### 2.13 Systemic administration of KYNA-1 to the experimental animals

For systemic chronic administration of KYNA-1, the SHR-24 transgenic rats were divided into placebo and KYNA-1 administrated groups (8 animals in each group, 4M/4F, bodyweight 250 ± 50 g). KYNA-1 (200mg kg^-1^) was administrated three times per week for three months. Sterile sodium chloride physiological solution was administrated to animals in the placebo group.

### 2.14 Sample preparation for determination of KYNA-1 and KP metabolites concentrations

Plasma samples for quantification were prepared as follows. First, 10 µL of internal standards 65 µL of water and 25 µL of TCA was added to 100 µL of plasma. Samples were vortexed and centrifuged at 30 000 g for 10 min., at 10 °C. Supernatants were then transferred to vials (Waters, Milford, MA, USA) and subjected to LC-MS analysis. Approximately 100 mg of brain tissue was weighed and homogenized in water (300 µL for 100 mg) using stainless steel beads (SSB16, Next Advance, USA) in FastPrep-24™ (MP Biomedicals, USA) at a vibrating speed 4.5 m s^-1^for the 20 s and centrifugated at 30 000 g for 10 min., at 10 °C. 200 µL of supernatant was transferred into a new tube, internal standard, and 600 µL of LC-MS acetonitrile (Honeywell, North Carolina, USA) was added for protein precipitation. After centrifugation, 700 µL of supernatant was evaporated using Savant Speed Vac SPD111V (ThermoFisher Scientific, USA). The samples were reconstituted in 50 µL of starting mobile phase, centrifugated, transferred into vials, and subjected to LC-MS analysis. Samples from the *in vitro*BBB permeability experiments were diluted 1:16 with starting mobile phase, centrifuged, transferred into vials, and subjected to LC-MS analysis.

### 2.15 Analysis of KYNA-1 and KP metabolites using liquid chromatography coupled to tandem mass spectrometry (UHPLC-MS/MS)

The concentrations of KYNA-1 and kynurenines were determined by ACQUITY UPLC I-Class coupled with triple quadrupole mass spectrometer ACQUITY XEVO TQD (Waters, Milford, MA, USA) with electrospray ionization (ESI) source in positive mode. MS/MS analysis was performed in the multiple reaction monitoring (MRM) mode. The electrospray capillary voltage was set to 2.5 kV; desolvation gas flow was 800 L/hr with a temperature of 350 °C and the source temperature was set to 150 °C. MassLynx software version 4.1 (Waters, Milford, MA, USA) was used for the data acquisition, and TargetLynx XS (Waters, Milford, MA, USA) was used for processing. Chromatographic separation was carried out on ACQUITY UPLC HSS T3 (100 × 2.1 mm, 1.8 µm particles) column (Waters, Milford, MA, USA) with gradient elution. Mobile phase A consisted of formic acid (Sigma-Aldrich, St. Louis, MO, USA) in LC-MS water (0.5%), and mobile phase B consisted of 100 % acetonitrile. The elution started at 5 % B (0 – 0.5 min.) gradually increasing to 95 % B (0.5 – 4.0 min.) held on 95 % B (4.0 – 4.5 min), then returning to 5 % and eventually re-equilibrating from 4.6 to 7.0 min. The column temperature was set to 40 °C, the flow rate was 0.5 ml min^-1^and the injection volume was 10 µL. The validation of the UHPLC-MS/MS method was carried out in compliance with the US Food and Drug Administration (FDA) [29] guideline. A complex overview of validation parameters is summarized in Supporting Information (Table S1 and Table S2).

### 2.16 MALDI imaging

KYNA-1 was administrated to SD rats. After 30 min rats were deeply anesthetized with tiletamine-zolazepam/xylazine anesthesia and perfused intracardially with phosphate buffer saline (PBS, 137 mM NaCl, 2.7 mM KCl, 10 mM Na_2_HPO_4_, 2 mM KH_2_PO_4_, pH 7.4) with heparin. Brain samples were cut coronally, frozen in 2-methyl butane (Sigma-Aldrich, St. Louis, MO, USA) over the fumes of liquid nitrogen, and stored at -80°C. 12-μm-thick brain sections were cut on a cryomicrotome (Leica CM 1850, Leica, Wetzlar, Germany) and thaw-mounted on precooled ITO glass slides (Bruker Daltonics, Bremen, Germany). Matrix deposition was carried out using an Ace vacuum sublimation apparatus (Sigma-Aldrich, St. Louis, MO, USA). The sublimation chamber was placed on a sand bath on a hot plate with temperature regulation (Magnetic Stirrer MR Hei-Tec) set to 140°C. Dried glass slides with brain tissue were attached to the inner condensation top. The top was filled with ice slush and the temperature was left to settle for 10 min. 300mg of 2,5-dihydroxybenzoic acid (DHB) powder (Sigma-Aldrich, St. Louis, MO, USA) was put on the outer bottom part and the apparatus was sealed and evacuated using hybrid pump RC 6 (Vacuubrand, Germany) at sublimation conditions: 140°C, 50 mbar, 10 min. MALDI imaging was performed using Bruker UltrafleXtreme MALDI TOF/TOF instrument equipped with a 2 kHz laser Smartbeam II (Bruker Daltonics, Bremen, Germany) operating in reflector mode, in both positive and negative mode with a mass range 400 – 2000 m/z. The laser spot size was set at small focus with a laser intensity of 70 % and 200 laser shots per pixel. Red phosphorus (suspension in acetonitrile; 10 mg ml^-1^, Sigma-Aldrich, St. Louis, MO, USA) was used as a calibrant. Images were acquired at a spatial resolution of 75 µm. Data were acquired and analyzed using FlexControl v3.4 and FlexImaging v4.1 software (Bruker Daltonics, Bremen, Germany).

### 2.17 Simoa digital ELISA

Concentrations of total tau proteins in rat plasma samples were measured by Simoa digital ELISA, using an HD-1 Analyzer (Quanterix, Boston, USA). For the analysis, the Simoa™ Mouse TAU Discovery kit was used (Quanterix, Boston, USA). Briefly, rat plasma samples were thawed, centrifuged for 10 min at 25 °C, and 20 000 x g. Supernatants were transferred onto Simoa sample plates together with a tau calibrator (part of the kit). Each measurement was done in duplicate according to the manufacturer’s recommendations. Concentrations were calculated using Simoa™ HD-1 instrument software (Quanterix, Boston, USA).

### 2.18 Immunocytochemical staining

Primary rat glial cells (astrocytes and microglia) and neuroblastoma cells were fixed for 15 min in 4% paraformaldehyde (Sigma-Aldrich, St. Louis, MO, USA) and washed with PBS. Cells were blocked for 60 min in 5% BSA in PBS. Sections were incubated overnight with primary antibodies: polyclonal rabbit anti-rat GFAP antibody (Abcam, Cambridge, UK), monoclonal mouse anti-CD16 antibody (Santa Cruz Biotechnology, Texas, USA), monoclonal mouse anti-CD32 antibody (Santa Cruz Biotechnology, Texas, USA), monoclonal mouse anti-CD163 antibody (Abcam, Cambridge, UK), polyclonal rabbit anti-Iba-1 antibody (Wako Chemicals USA, Inc.), polyclonal rabbit anti-GFAP antibody (Abcam, Cambridge, UK) and monoclonal mouse anti-tau antibody (DC190, epitope aa368-376, Axon-Neuroscience R&D, Bratislava, Slovakia). After washing, the sections were incubated for 1 hour in goat anti-rabbit or goat anti-mouse AlexaFluor488/546 secondary antibodies (Invitrogen Life Technologies, Carlsbad, CA). Sections were mounted and examined using an LSM 710 confocal microscope (Zeiss, Jena, Germany).

### 2.19 Immunohistochemistry

Animals were anesthetized by tiletamine-zolazepam/xylazine anesthesia and perfused intracardially with PBS with heparin. The brain was removed and embedded in a cryostat embedding medium (Leica, Wetzlar, Germany) and frozen above the surface of liquid nitrogen. 12-μm-thick brain sections were cut on a cryomicrotome, fixed onto poly-L-lysine coated slides, and left to dry at room temperature for 1 hour. Sections were fixed for 15 min in 4% paraformaldehyde and blocked for 60 min in blocking solution (DAKO, Mississauga, Ontario, Canada). Sections were incubated overnight in polyclonal rabbit anti-GFAP primary antibody and monoclonal mouse anti-tau antibody (DC190). After washing, the sections were incubated for 1 hour in goat anti-rabbit or goat anti-mouse AlexaFluor488/546 secondary antibodies. Brain slices were mounted (Vector Laboratories, Burlingame, CA, USA) and examined using an LSM 710 confocal microscope.

### 2.20 Western blot

Cells and brain tissue were homogenized into lysis buffer (200 mM Tris, pH 7.4, 150 mM NaCl, 1 mM EDTA, 1 mM Na_3_VO_4_, 20 mM NaF, 0.5% Triton X-100, 1 × protease inhibitors complete EDTA-free from Roche, Mannheim, Germany). Total protein concentration of prepared cell extracts was measured by Bio-Rad protein assay (Bio-Rad Laboratories GmbH, Colbe, Germany). A total of 20 μg of the proteins was separated onto 12% SDS-polyacrylamide gels and transferred to a nitrocellulose membrane in 10 mM N-cyclohexyl-3-aminopropanesulfonic acid (CAPS, pH 11, Roth, Karlsruhe, Germany). The membranes were blocked in 5% milk in Tris-buffered saline with Tween 20 (TBS-T, 137 mM NaCl, 20 mM Tris-base, pH 7.4, 0.1% Tween 20 all from Sigma-Aldrich) for 1 hour and incubated with rabbit polyclonal anti-GFAP primary antibody, monoclonal mouse anti-pSer202/pThr205 (Thermo Fisher Scientific, Massachusetts, USA), monoclonal mouse anti-pThr181 (Thermo Fisher Scientific, Massachusetts, USA), polyclonal rabbit anti-pThr212 (Thermo Fisher Scientific, Massachusetts, USA) overnight at 4°C. Membranes were incubated with horseradish peroxidase (HRP)-conjugated secondary antibody in TBS-T (Dako, Glostrup, Denmark) for 1 hour at room temperature. Immunoreactive proteins were detected by chemiluminescence (SuperSignal West Pico Chemiluminescent Substrate, Thermo Scientific, Pittsburgh, USA) and the signals were digitized by Image Reader LAS-3000 (FUJIFILM, Bratislava, Slovakia).

### 2.21 Data analysis

Relative staining patterns and intensity of projections were visualized by confocal microscopy and Image Reader LAS-3000 and evaluated by image analysis. ImageJ was used for the evaluation and quantification of chemiluminescent signal, immunohistochemical and immunocytochemical slides. We quantified 10 slices from each sample. For semiquantitative analysis, the colour pictures were converted to grayscale 8-bit TIFF file format, and regions of interest were analyzed with ImageJ software. The grayscale 8-bit images were converted to thresholded 1-bit images, on which the number of immunolabeled structures localized in the area of interest was measured. Then the average intensity/pixel values of each area were calculated, and the average intensity/pixel values representing the background intensity were subtracted from those of immunolabeled areas. All results were expressed as mean ± standard deviation (SD). Differences between means were analyzed with an independent two-tailed Student’s t-test and one-way ANOVA (Prism 6.0 software, GraphPad, Inc., SanDiego, CA). Differences at p<0.05 were regarded as statistically significant.

All data analysis and presentation techniques adhere to the BJP Editorial checklists.

## 3 RESULTS

### 3.1 KYNA-1 toxicity

KYNA-1 is the amide of KA (Figure 1a). To determine the cytotoxicity of KYNA-1 a series of KYNA-1 dilutions (from 500 mM to 100 nM) was added to primary rat astrocytes and neuroblastoma cells. 24 hours post-treatment, cell toxicity was measured and IC_50_values were calculated. Results from the assay showed low cytotoxicity. The IC_50_values were 16.1 ± 0.9 mM in primary cultures of rat astrocytes and 15.9 ± 1.49 mM in neuroblastoma cells (Figure 1b and c).

**Figure 1.**
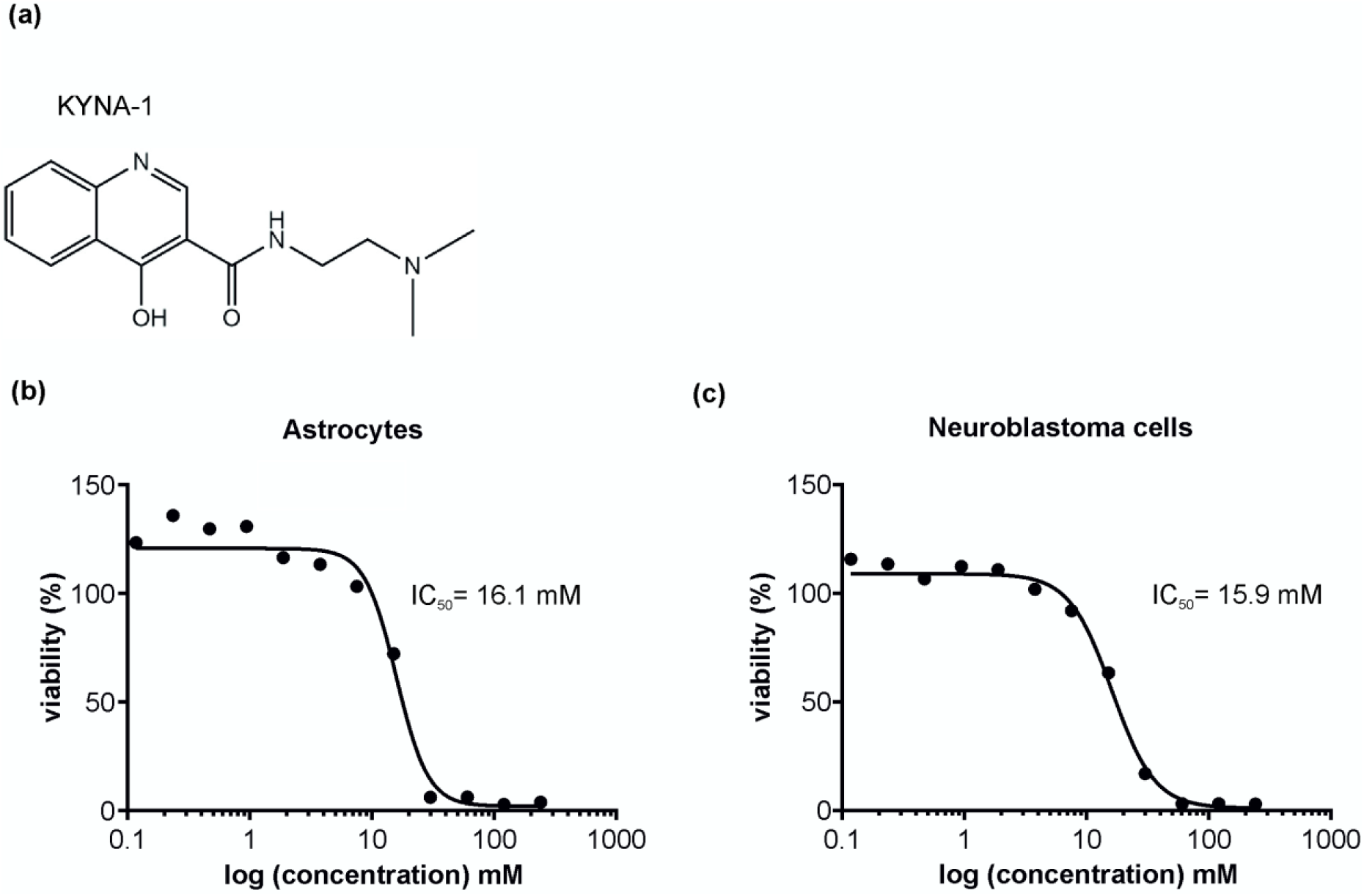
KYNA-1 cytotoxicity. Chemical formula of KYNA-1 analog (a). Cytotoxicity assay was performed after 24 hours of KYNA-1 treatment to assess its effect on viability of primary rat astrocytes (b) and human neuroblastoma cells (c). The half-maximal inhibitory concentration (IC50) at which KYNA-1 exerts 50% of its inhibitory effect was calculated from the concentration-response curve. Data shown as means ±SD (n=9).

### 3.2 KYNA-1 blood-brain barrier permeability

To examine whether the KYNA-1 analog could penetrate across BBB, we used an *in vitro*BBB model (Figure 2a– schema). In brief, primary rat endothelial cells were seeded onto the apical side of Transwell inserts and co-cultured with primary rat astrocytes for 7 days to encourage the formation of a tight cell monolayer expressing a tight junction proteins. Immunocytochemical staining confirmed that primary rat endothelial cells expressed the tight junction proteins zonula occludens-1 (ZO-1) and claudin-5 (Figure 2b). Isolated primary rat astrocytes showed high expression of GFAP protein (Figure 2b). Expression of tight junction proteins in our BBB model reflects the protein-related properties of *in vivo*BBB. These results indicated that the *in vitro*BBB model can be used for permeability experiments.

**Figure 2.**
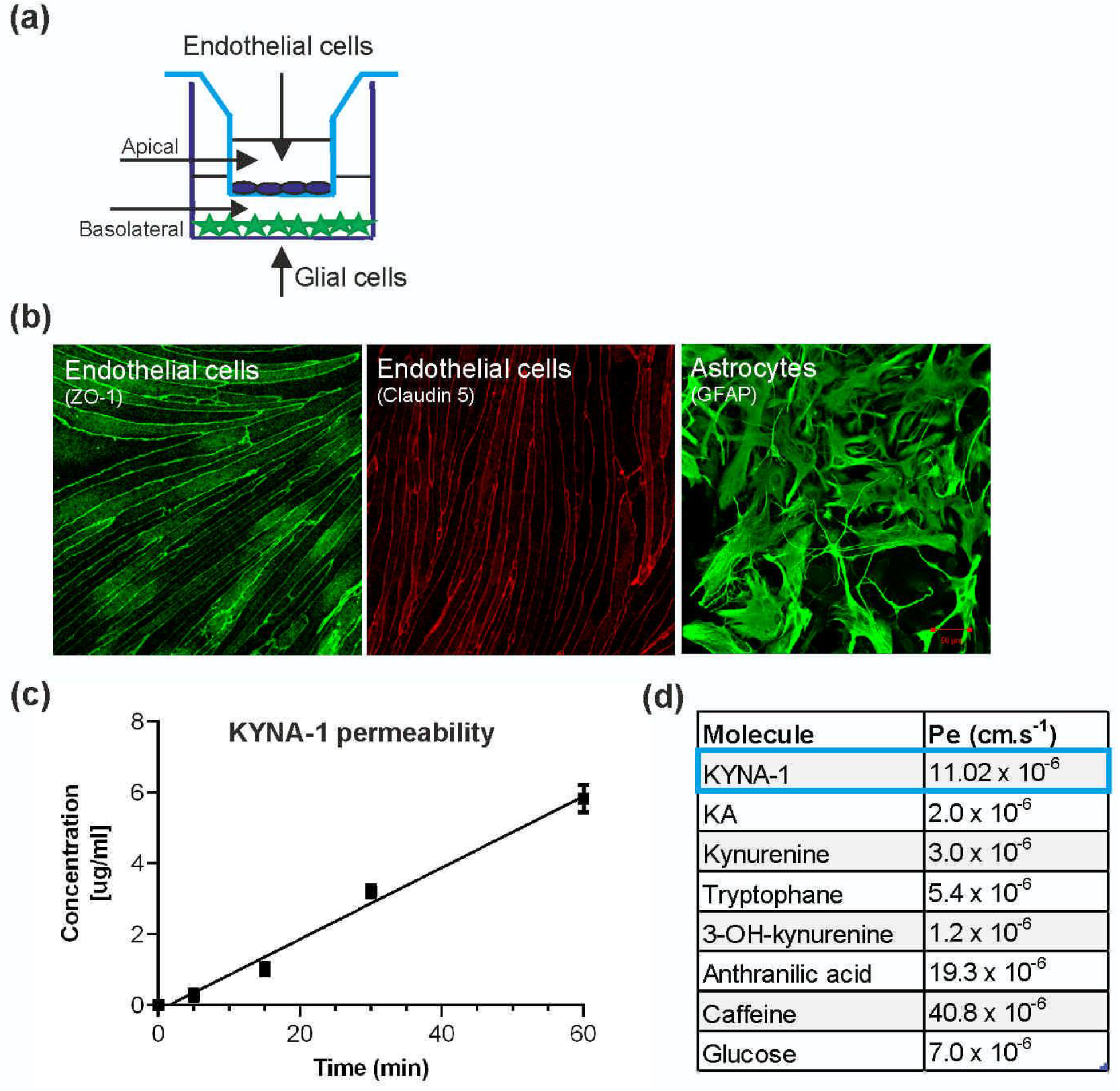
KYNA-1 permeability across the in-vitro blood-brain barrier model. Endothelial cells are seeded onto porous inserts (apical compartment) and mixed glial cells are at the bottom of the plate (basolateral compartment) (a). The rat brain endothelial cells showed typical spindle-shaped morphology and expressed tight junction proteins ZO-1 and claudin-5 (b). Penetration of KYNA-1 analog through the in vitro BBB model. Permeability of KYNA-1 increases in time-dependent manner (c). Permeability coefficient of KYNA-1 compared to other kynurenines, caffeine and glucose [46] (d).

KYNA-1 analog (final concentration 3.8 µM) was added to the luminal part of the BBB model and at different time points (5, 15, 30, and 60 min.) aliquots of media were collected from the abluminal chamber. The concentration of KYNA-1 was quantified using LC-MS/MS method. The results showed that KYNA-1 was able to cross the endothelial monolayer (Figure 2c). Its permeability significantly increased in a time-dependent manner. The endothelial permeability coefficient (Pe) of KYNA-1 was calculated 1.1 × 10^−7^cm s^-1^. The Pe of KYNA-1 was 5.5 times higher than Pe of KA (Figure 2d).

### 3.3 Plasma and brain tissue pharmacokinetic profile of KYNA-1 after single and chronic i.p. administration

After the single intraperitoneal dose (200 mg.kg^-1^) the levels of KYNA-1 peaked in serum and brain tissue at 30 minutes and then slowly decreased with a half-life of 2.6 h (Figure 3a). The calculated pharmacokinetic parameters are in Table 1. The AUC_0-t_value was 191507.32 ng ml^-1^h^-1^. The maximum brain levels were approximately 45-53 times lower than in the plasma (Figure 3b). The unidirectional influx rate of KYNA-1 in the frontal cortex and brainstem was 11.85±1.6 µl g^-1^min^-1^and 7.05±1.9 µl g^-1^min^-1^respectively (Figure 3c).

**Figure 3.**
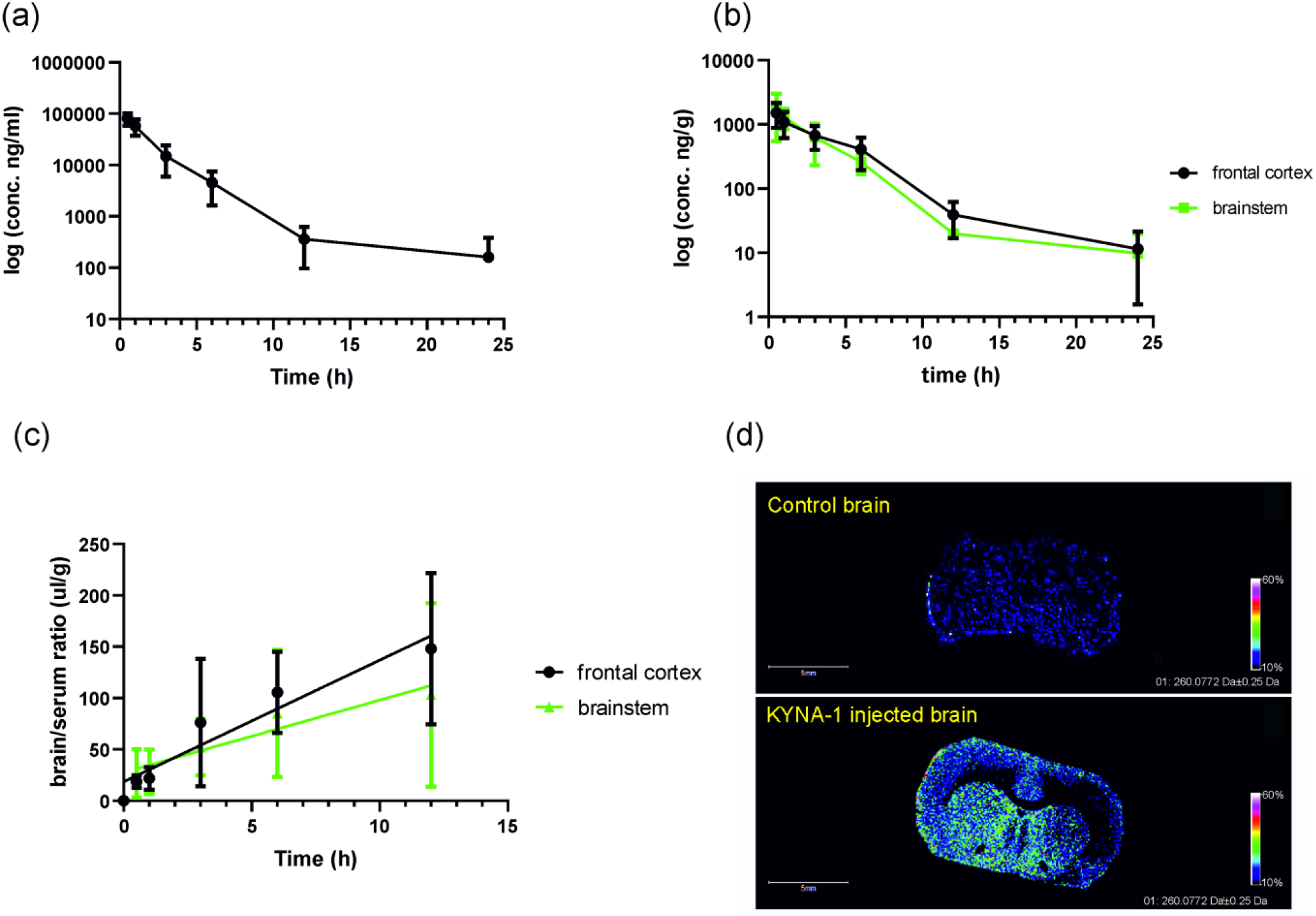
Concentration of KYNA-1 in plasma (a) and brain tissue (b) after single intraperitoneal injection (200mg/kg). The brain/serum ratio of KYNA-1 (c). MALDI imaging showed the distribution of KYNA-1 (m/z 260.07) in coronal section of brain tissue (d). Data shown as means ±SD (n=8 animals).

**Table 1.**
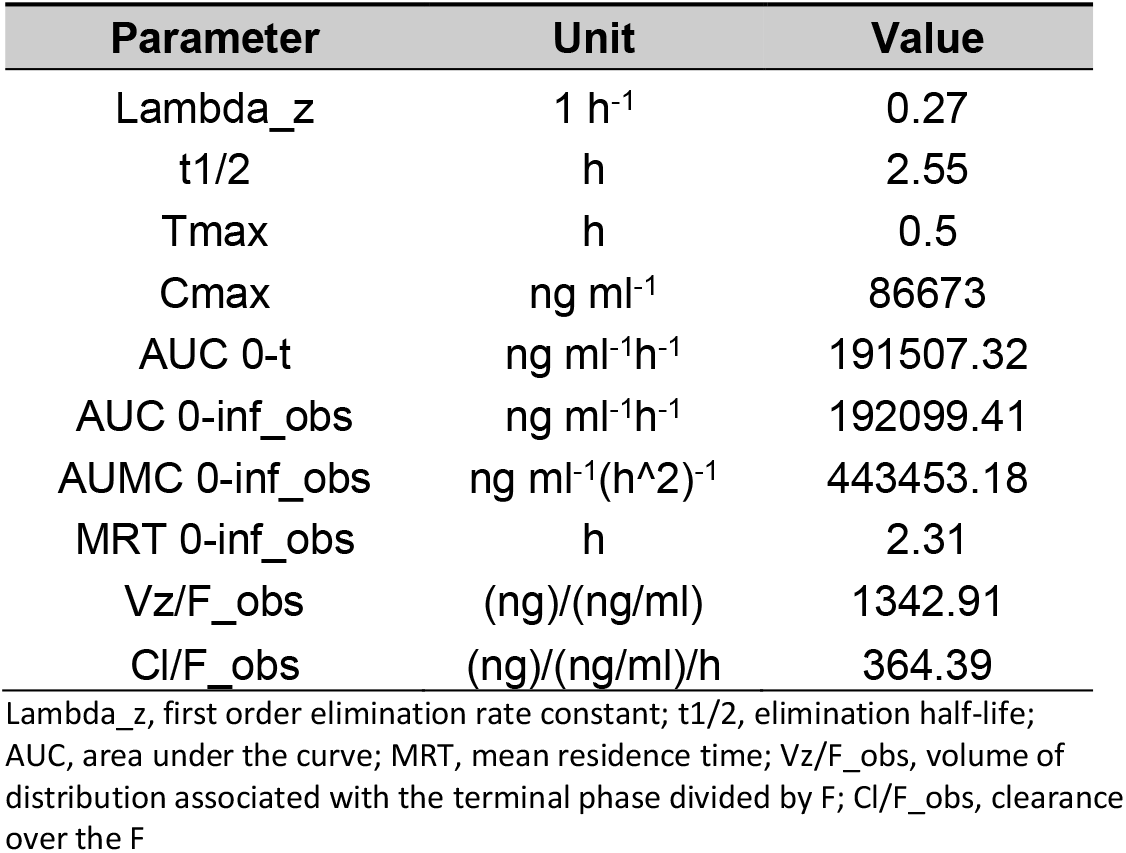
Pharmacokinetic parameters after a single intraperitoneal administration of 200mg/kg KYNA-1 to SD rats.

| Parameter | Unit | Value |
| --- | --- | --- |
| Lambda<sub>z</sub> | 1 h<sup>-1</sup> | 0.27 |
| t<sub>1/2</sub> | h | 2.55 |
| T<sub>max</sub> | h | 0.5 |
| C<sub>max</sub> | ng ml<sup>-1</sup> | 86673 |
| AUC 0-t | ng ml<sup>-1</sup>h<sup>-1</sup> | 191507.32 |
| AUC 0-inf_obs | ng ml<sup>-1</sup>h<sup>-1</sup> | 192099.41 |
| AUMC 0-inf_obs | ng ml<sup>-1</sup>(h<sup>2</sup>)<sup>-1</sup> | 443453.18 |
| MRT 0-inf_obs | h | 2.31 |
| V<sub>z</sub>/F<sub>obs</sub> | (ng)/(ng/ml) | 1342.91 |
| Cl/F<sub>obs</sub> | (ng)/(ng/ml)/h | 364.39 |

Lambda<sub>z</sub>, first order elimination rate constant; t<sub>1/2</sub>, elimination half-life; AUC, area under the curve; MRT, mean residence time; V<sub>z</sub>/F<sub>obs</sub>, volume of distribution associated with the terminal phase divided by F; Cl/F<sub>obs</sub>, clearance over the F

By MALDI-imaging we showed the distribution of KYNA-1 from the periphery into the brain. High-resolution images confirmed the presence of KYNA-1(m/z 260.08) in rat coronal brain sections 30 min after the administration. KYNA-1 showed diffuse signal in all brain areas without more significant accumulation (Figure 3d).

Next, we analyzed the levels of KP metabolites (kynurenine, kynurenic acid, anthranilic acid, and xanthurenic acid) in the plasma and brain tissue of KYNA-1 injected animals. We found that 30 minutes after administration of KYNA-1 the plasma levels of KA significantly increased in comparison to basal levels (6.51±1.32ng ml^-1^at t0 vs 475.4±188.5ng.ml^-1^at t30), then returning to basal values after 12 hours. The plasma levels of anthranilic acid gradually increased up to 12 hours, then slowly returned to basal values at 48 hours. The levels of kynurenine and xanthurenic acid remained unchanged (Figure 4a). Interestingly, we observed no change of kynurenic acid levels in the brainstem and frontal cortex at any time point. The brain tissue levels of other kynurenine metabolites remained unchanged as well (Figure 4b).

**Figure 4.**
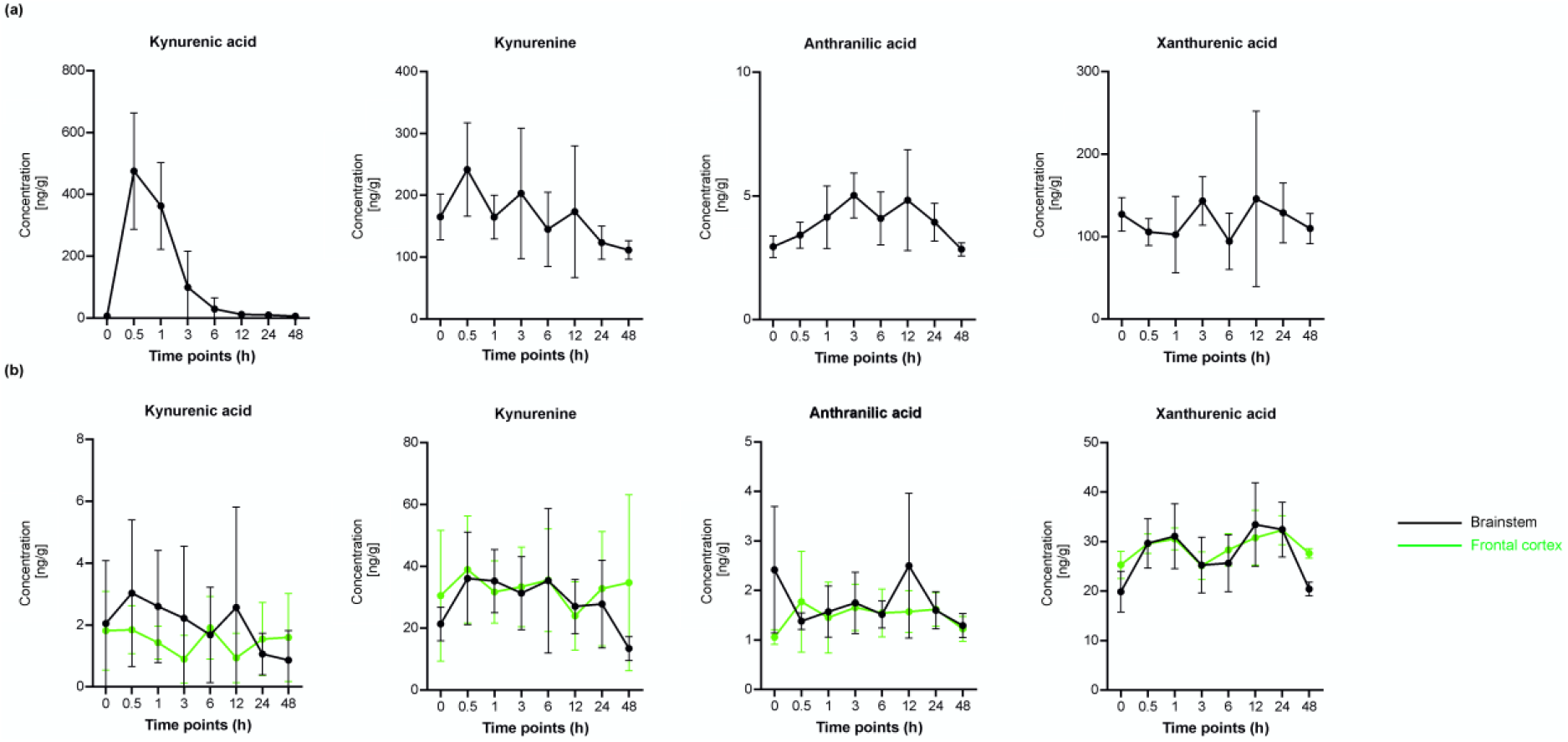
Concentrations of KP metabolites in plasma (a) and brain tissue (brainstem and cortex) (b) measured by UHPLC-MS/MS after KYNA-1 administration. Data are shown as mean ± SD (n=8 per each time

During the chronic administration, KYNA-1 was injected into the transgenic rat model for tauopathy (SHR-24) three times per week for three months. In the placebo group, phosphate buffer was injected. After three months, the concentrations of KYNA-1 and KP metabolites were measured in plasma samples and brain tissue of placebo and KYNA-1 injected group by validated LC-MS/MS method. The plasma and brain concentrations of KYNA-1 analog and KP metabolites are presented in Table S3. The data showed significant decrease of kynurenic acid and xanthurenic acid in plasma of KYNA-1 treated animals. There was no significant difference in KP metabolites in brain tissue between placebo and KYNA-1 treated animals.

### 3.4 KYNA-1 modulate the level of sarkosyl-insoluble tau in brain tissue of transgenic animals

To examine whether KYNA-1 analog affected tau pathology, we analyzed total tau in plasma and brain tissue as well as levels of hyper-phosphorylated and aggregated forms of tau protein. Total tau was determined with an anti-tau antibody (DC190, recognizing epitope aa368-376). The soluble fraction of tau protein isolated from the brainstem contained two major bands; truncated tau protein (aa151-391/3R, 35 kDa) and endogenous full-length tau (aa1-441/4R, 50 kDa) (Figure 5a). In KYNA-1 treated animals, we found no difference in the expression of soluble tau compared with placebo group (Figure 5b).

**Figure 5.**
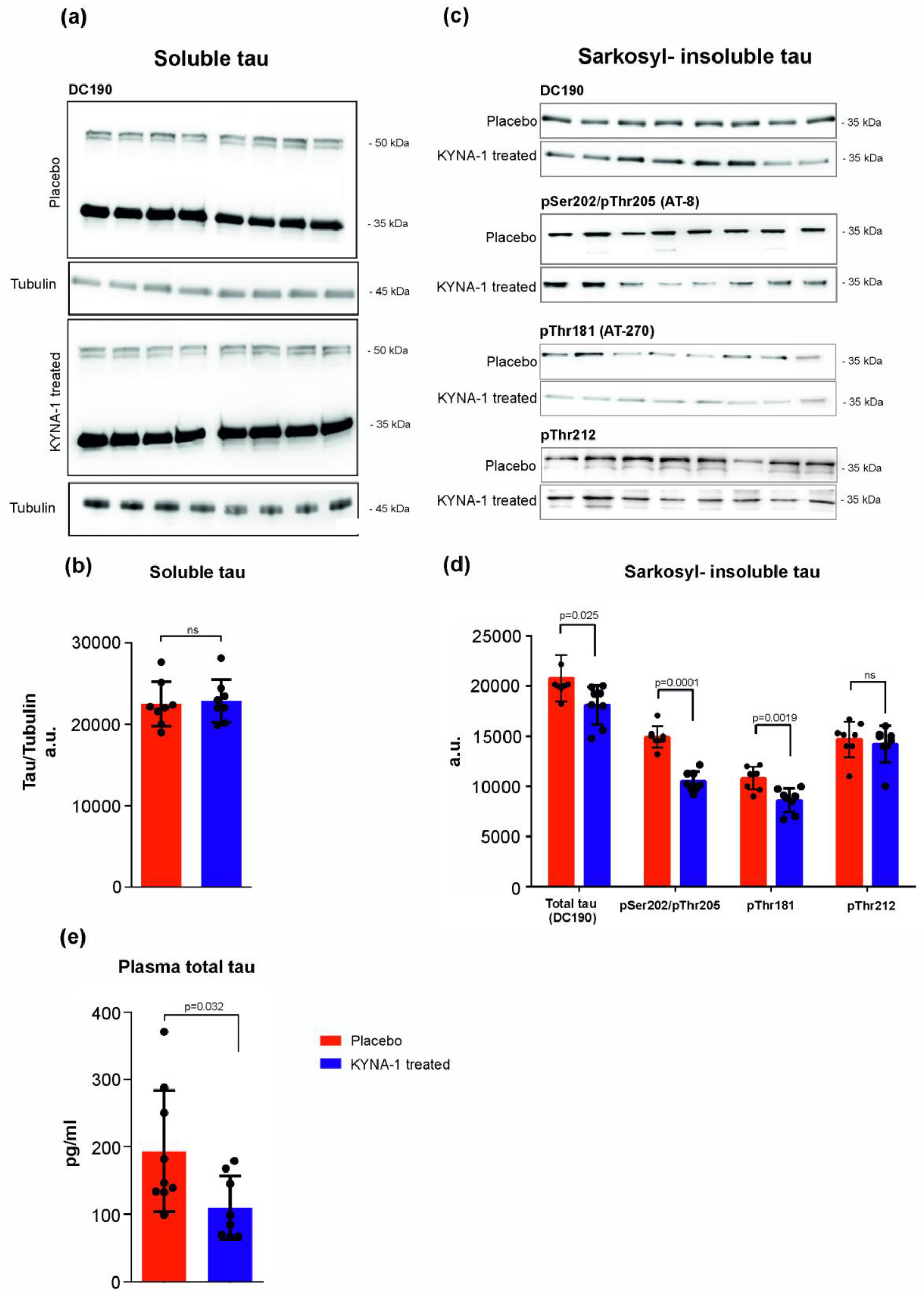
Effect of KYNA-1 on neurofibrillary pathology in transgenic rat model for tauopathy SHR-24. Western blot analysis of soluble tau protein. Representative cropped images of the membranes for tau staining and tubulin are shown. Tubulin was used as a loading control (a). Quantification of soluble tau protein showed no changes between KYNA-1 treated and placebo group. The data are shown as mean ± SD (n= 8) (b). Biochemical analysis of sarkosyl-insoluble tau. The membranes were stainned against three diferrent phospho-dependent tau epitopes (pSer202/pThr205, pThr181, pThr212) and total tau protein (DC190 anti-tau antibody) (c). Quantification of tau proteins showed reduction of sarkosyl-insoluble tau after KYNA-1 treatment compare to control animals. The data are shown as mean ± SD (n= 8) (d). The amount of total tau protein was analysed in plasma by digital ELISA (Simoa HD-1 analyser). The results showed significant reduction of tau protein in plasma after KYNA-1 treatment. The data are shown as mean ± SD (n= 8) (e).

The presence of sarkosyl-insoluble tau protein in brain tissue is the main pathological feature of tauopathies. Therefore, we analyzed sarkosyl-insoluble protein extracts from the brainstem of SHR-24 transgenic rats after KYNA-1 treatment. To monitor the level of phosphorylation of insoluble tau protein, we performed biochemical western blot analysis using specific phospho-tau antibodies against pThr181, pSer202/pThr205, and pThr212 (Figure 5c). The results showed that the total amount of sarkosyl-insoluble tau stained with DC190 antibody significantly decreased in KYNA-1 treated animals compared to the placebo group (Figure 5d). Besides that, we found significantly lower levels of pSer202/pThr205 (placebo: 14 090 ± 2340; KYNA: 7889 ± 3160; n= 8, p< 0.001) and pThr181 positive tau (placebo: 14 886 ± 3056; KYNA: 10289 ± 4209; n= 8, p< 0.038) in KYNA-1 treated rats compared to placebo. There was no change in amount of pThr212 positive sarkosyl-insoluble tau (placebo: 17 114 ± 1048; KYNA-1: 16178 ± 1224; n= 8, p< 0.098, Figure 5d).

We further analyzed the total tau protein in plasma by supersensitive digital ELISA. Results showed that treatment with KYNA-1 decreased the levels of total tau protein in plasma (placebo: 249.5 ± 52.08; KYNA-1: 109.6 ± 16.72; p= 0.02; n= 8, Figure 5e).

Altogether the above data suggest that KYNA-1 reduced hyperphosphorylated aggregated tau in the brain tissue of transgenic rats as well as soluble plasma tau.

### 3.5 Effect of KYNA-1 on GFAP expression *in vivo*in a transgenic rat model for tauopathy

Neuroinflammation is an important hallmark of tauopathies. Activation of astrocytes closely correlates with neurofibrillary pathology. In this experiment, we asessed how KYNA-1 analog affects the activity of astrocytes in brain tissue with tau pathology. Immunohistochemistry staining of brain tissue from SHR-24 animals revealed reactive astrocyte phenotype with intense GFAP signal and highly ramified morphology with hypertrophic processes. In contrast, astrocytes in SHR rats showed stellate morphology and a low level of GFAP immunoreactivity (Figure 6a). We next semiquantified the GFAP expression in KYNA-1 treated animals. Compared to placebo group, KYNA-1 was able to modulate astrogliosis in brain area positive for neurofibrillary pathology and decrease GFAP reactivity in SHR-24 transgenic rats (placebo: 7876 ± 676.2; KYNA-1: 5249 ± 649.1; n=8; p= 0.0135) (Figure 6b and c).

**Figure 6.**
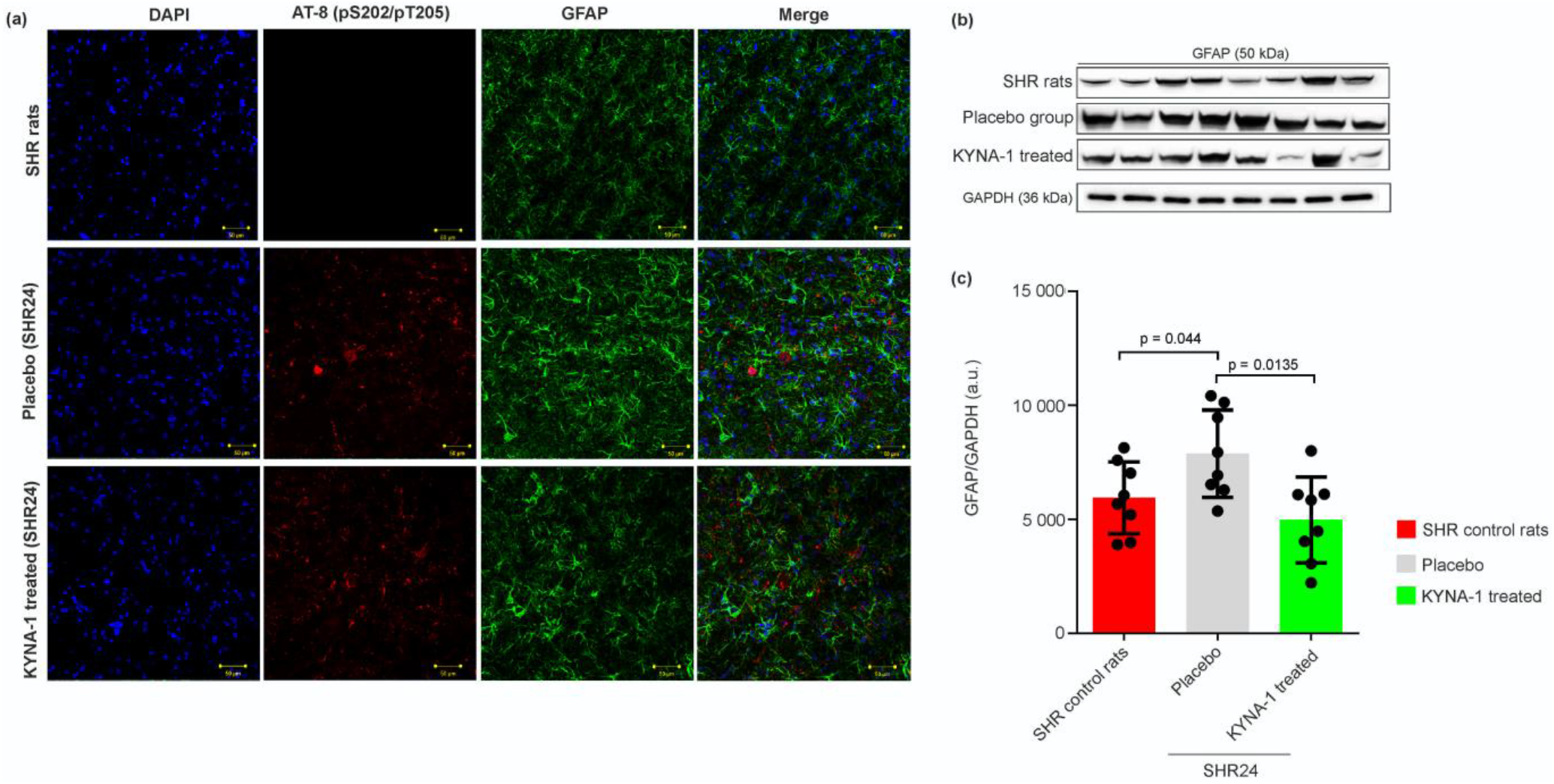
Effect of KYNA-1 on GFAP immunoreactivity of astrocytes in transgenic rat model for tauopathies. Immunohistochemical staining of astrocytes in control SHR rats and SHR-24 treated group. Rat astrocytes were stained for GFAP (green), tau neurofibrillary pathology was by anti-tau antibody pS202/pT205 (red). DAPI nuclear staining is shown in blue (a). The GFAP expression was analyzed at the protein level by western blot analysis, representative cropped images of the membranes for the GFAP and GAPDH as a loading control are shown (b). Semi-quantitative analysis of GFAP expression in placebo and KYNA-1 treated group compare to control SHR rats (c). The data are presented as the mean ± SD (n= 9).

KYNA-1 has the potential in reducing the glial activation associated with tau pathology.

### 3.6 Effect of KYNA-1 on LPS-induced stimulation of glial cells

We investigated the effect of KYNA-1 on LPS-induced cell toxicity and astrocyte activation. Primary rat astrocytes were incubated with KYNA-1 (final concentration 100 nM) for 1 hour and stimulated with LPS (500 ng.ml^-1^) for additional 23 hours (Figure 7a). Viability assay showed that LPS was able to induce significant cell death (control cells: 99.78 ± 2.41%; LPS: 161.9 ± 4.18%; p< 0.0001; n= 10). However, when KYNA-1 was added to LPS-stimulated cells toxicity significantly decreased (LPS: 161.9 ± 4.18%; LPS+ KYNA-1: 135.8 ± 7.1%; p= 0.0053; n= 10). KA showed similar effect (LPS: 161.9 ± 4.18%; LPS+ KA: 114.9 ± 3.4%; p= 0.0001; n= 10 (Figure 7b).

**Figure 7.**
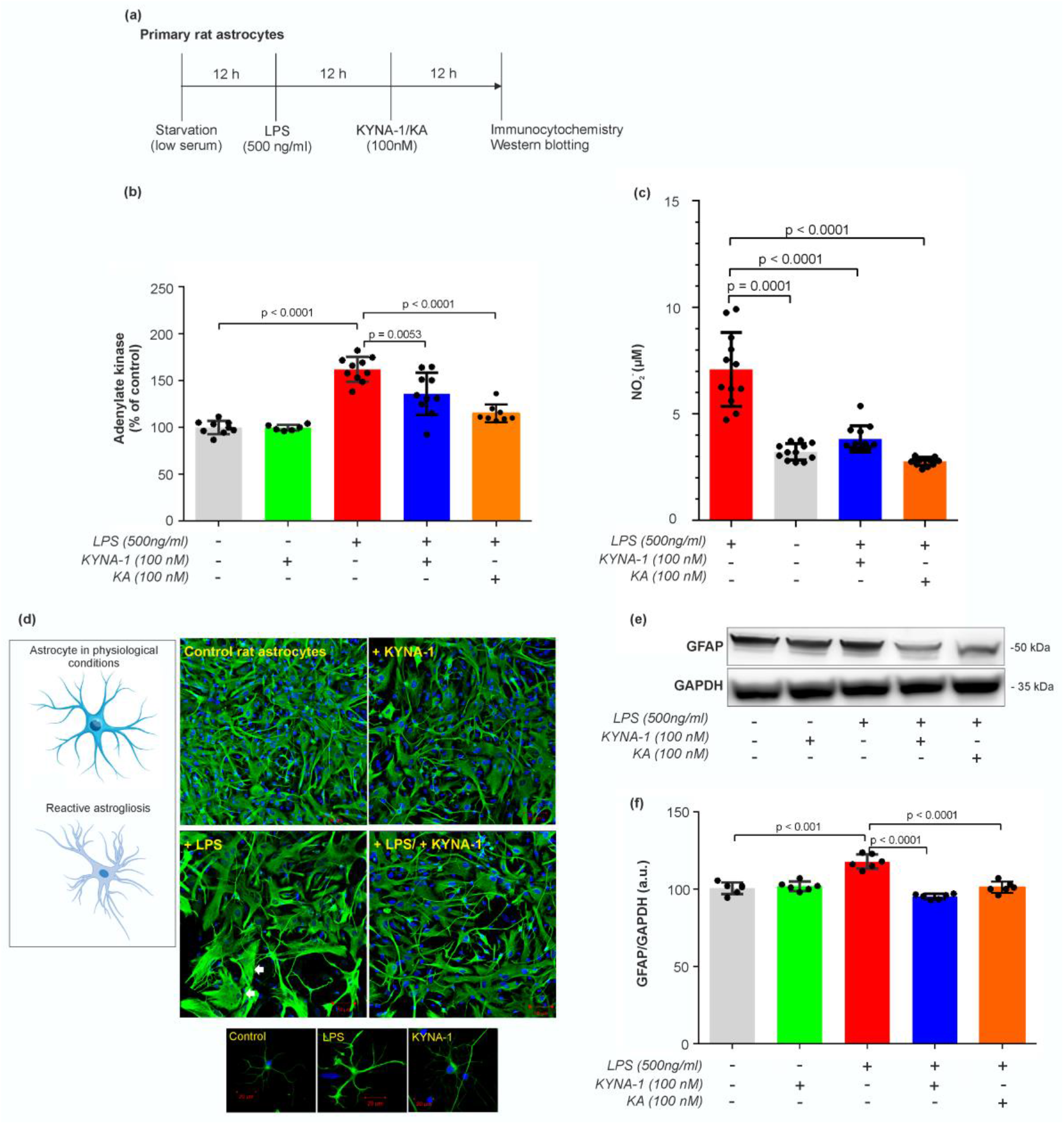
Effect of KYNA-1 on LPS-induced cell toxicity and stimulation of primary rat astrocytes. Timeline of in vitro experiments (a). KYNA-1 showed positive effect on astrocyte viability after LPS exposure (b). Release of nitric oxide by LPS stimulated mixed glia was signifficantly reduced by pre-treatment with KYNA-1 or KA (c). Immunocytochemical staining of astrocytes exposed to control medium, LPS and KYNA-1 analog. Rat astrocytes were stained for GFAP (green). DAPI nuclear staining is shown in blue. LPS induced reactive astrocyte phenotype. GFAP staining showed morphological changes from filamentous to flattened shape (white arrows) in response to LPS. KYNA-1 attenuated reactive phenotype of astrocytes (d). The GFAP expression was analyzed by western blot, representative cropped images of the membranes for the GFAP and GAPDH as a loading control are shown (e). GFAP expression levels decreased following KYNA-1 as well as KA stimulation (f). The data are presented as the mean ± SD (n=6).

Activation of glial cells is accompanied by the production of reactive nitrogen species. Thus, we studied the effect of KYNA-1 on NO production by astrocytes. The LPS significantly induced NO release (control: 3.2 ± 0.4 µM; LPS: 7.1 ± 2.7 µM; p= 0.0001; n= 12). Addition of KYNA-1 or KA significantly inhibited the NO production by cells (LPS: 7.1 ± 2.7 µM; LPS+KYNA-1: 3.8 ± 0.6 µM; LPS+KA: 2.8 ± 0.2 µM; p= 0.0001; n= 12) (Figure 7c).

We also analyzed the effect of KYNA-1 on GFAP expression. Immunofluorescence analysis of primary rat astrocytes showed stellate morphology in control conditions. Conversely, LPS-treated astrocytes exhibited higher GFAP signal and ramified morphology with hypertrophic processes(Figure 7d). Consistently, the expression of GFAP measured by western blot analysis was increased after LPS treatment (control: 100.6 ± 1.6; LPS: 114.8 ± 1.7; n= 6; p< 0.0001). The pretreatment with either KYNA-1 (LPS: 114.8 ± 1.7; KYNA-1 + LPS: 95.2 ± 0.7 n= 6; p< 0.0001) or KA (LPS: 114.8 ± 1.68; KA+ LPS: 101.2 ± 3.6 n= 6; p< 0.0001) markedly attenuated LPS-mediated upregulation of GFAP expression (Figure 7e and f).

Pathophysiological processes such as neuroinflammation and microglial activation are associated with multiple neurodegenerative diseases. Microglial heterogeneity is visible in their morphology and expression of cellular markers. Using the primary microglia cultures, we analyzed protein markers of the M1 (CD16 and CD32) and M2 (CD163) phenotypes after LPS stimulation. We found that KYNA-1 increased expression of CD163 marker (+LPS: 4025 ± 136.4; +LPS+KYNA-1: 6561 ± 692.7; n= 5; p= 0.0071) (Figure 8c and d). Furthermore, the expression of pro-inflammatory CD16 (+LPS: 2858 ± 155.5; +LPS+KYNA-1: 1236 ± 189.5; n= 5; p= 0.0002) and CD32 (+LPS: 2859 ± 551.9; +LPS+KYNA-1: 1214 ± 93.75; n= 5; p= 0.0148) markers was significantly decreased (Figure 8a, b and d).

**Figure 8.**
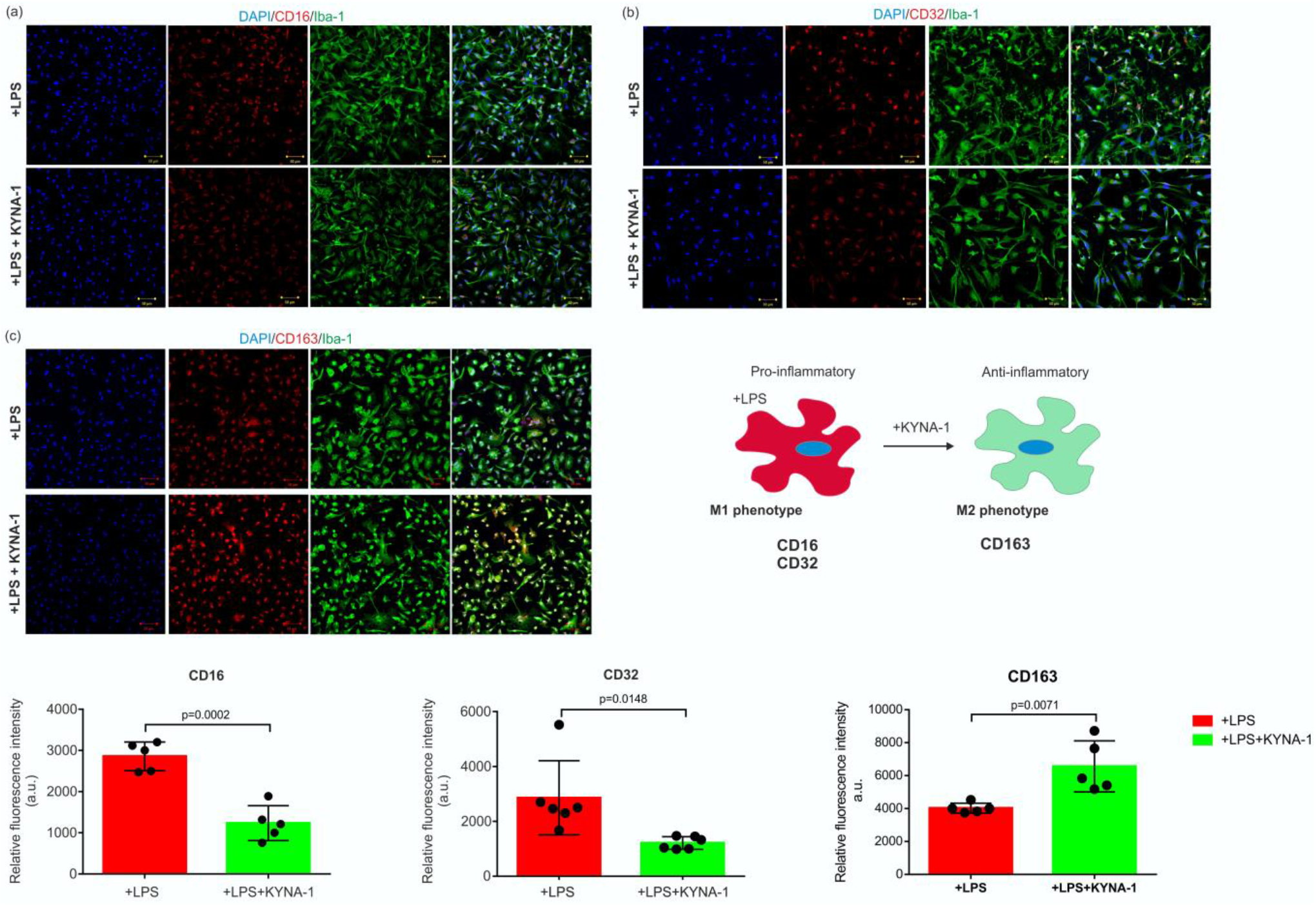
KYNA-1 treatment restores M2 phenotype in microglia activated by LPS. Immunocytochemical staining of Iba-1 positive primary rat microglia for CD16 marker (CD16-red; Iba-1-green). DAPI nuclear staining is shown in blue (a). Immunocytochemical staining of Iba-1 positive primary rat microglia for CD32 marker (CD32-red; Iba-1-green). DAPI nuclear staining is shown in blue (b). For detection of the M2 phenotype we used CD163 marker (CD163-red; Iba-1-green). DAPI nuclear staining is shown in blue (c). Quantification of M1 and M2 markers showed significant changes in LPS challenged microglia after KYNA-1 treatment. The data are representative of two experiments and are shown as mean ± SD (n= 6).

Together, these studies suggest that KYNA-1 can significantly decreases the activation of primary rat astrocytes during neuroinflammatory process. The results also showed that KYNA-1 stimulates anti-inflammatory, scavenging, and regenerative M2 phenotype in microglia.

### 3.7 Astrocytes in a multi-component cell model system modulate phosphorylation of truncated tau protein

Next, we used, the multi-component cellular model system (Figure 9a) for the study of molecular interactions between SH-SY5Y neuroblastoma cells expressing human truncated tau protein (aa 151-391/4R) and primary rat astrocytes. After five days SH-SY5Y neuroblastoma cells became fully matured into neuron-like phenotype. The cells were positive for neuron-specific and synaptic proteins such as synaptosomal-associated protein 25 (SNAP25, Supplementary Figure. S1a), beta III Tubulin (Supplementary Figure. S1b), Neurofilament SMI-312 (Supplementary Figure. S1c), and amphiphysin (Supplementary Figure. S1d). The results from immunocytochemical staining showed the re-distribution of these proteins from the cellular body into cell projections.

**Figure 9.**
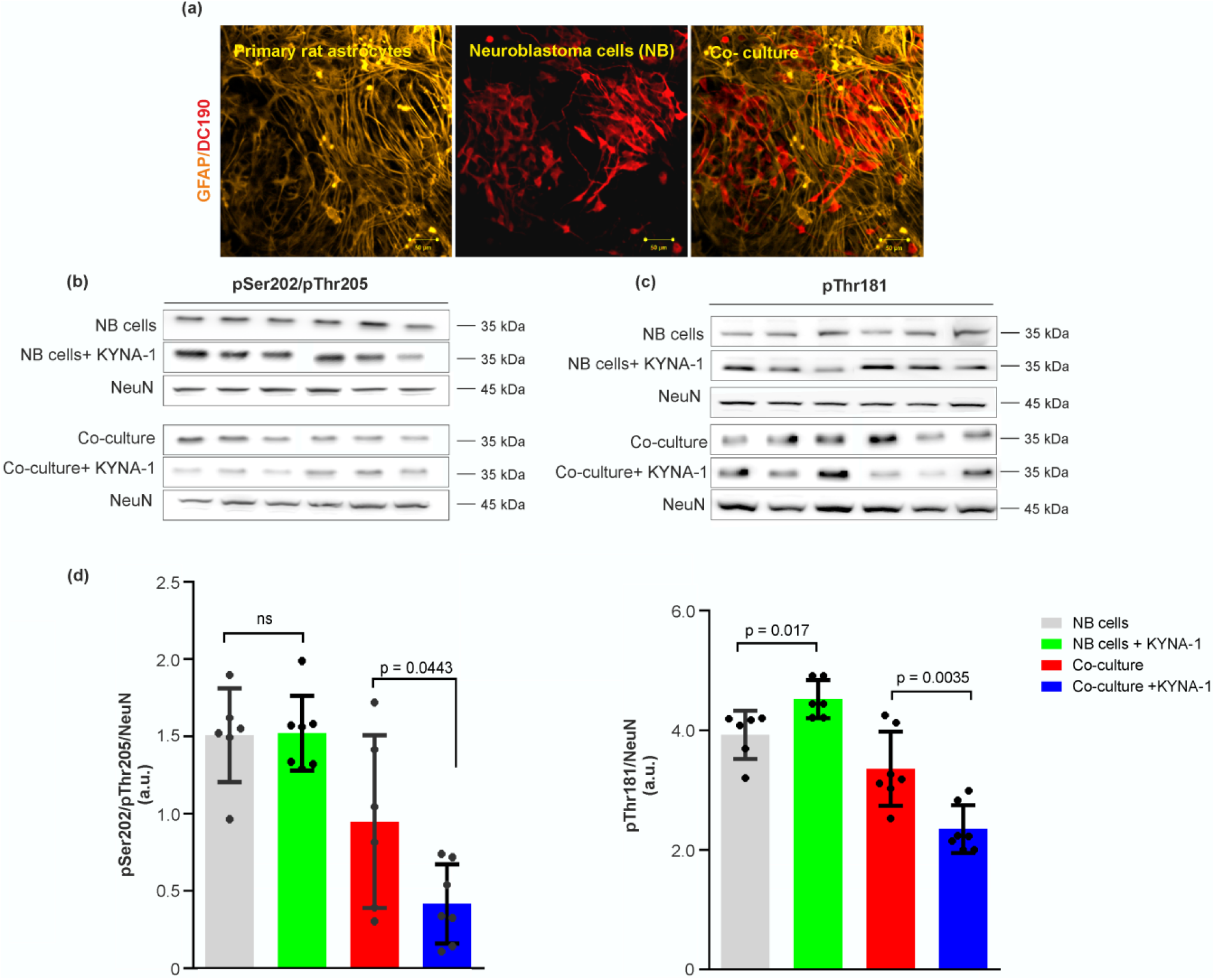
KYNA-1 showed effect on tau phosphorylation in multi-component cell system. Our multi-component cell system composed of primary rat astrocytes and neuroblastoma cells (SHSY-5Y) inducible expressing truncated tau protein (aa 151-391/4R). Immunocytochemical staining of astrocytes stained for GFAP (orange) and neuroblastoma cells stained for tau (DC190 anti-tau antibody, red) (a). Western blot analysis of pSer202/pThr205 tau phosphorylation in neuroblastoma cells and multi-component cell system. NeuN was used as a loading control of neuroblastoma cells (b). Western blot analysis of phosphorylated tau on pThr181 in neuroblastoma cells and multi-component cell system. NeuN was used as a loading control of neuroblastoma cells (c). Quantification of pSer202/pThr205 and pThr181 positive tau showed reduction of phosphorylation in multi-component cell system after KYNA-1 treatment (e). The data are shown as mean ± SD (n= 6).

Using our *in vitro*multi-component cell model and western blot analysis we studied changes of tau phosphorylation on several AD relevant epitopes: pSer202/pThr205, pThr181 in response to KYNA-1 treatment.

The amount of tau phosporylated on pSer202/pThr205 remain unchanged in KYNA-1 treated neuroblastoma cells (control: 1.5 ± 0.12, KYNA-1: 1.52 ± 0.09, n= 7). pThr181 phosporylation was slightly increased (control: 3.92 ± 0.16, KYNA-1: 4.31 ± 0.13, n= 7, p=0.017). Interestingly, when neuroblastoma cells were co-cultivated with astrocytes, the levels of both phosporylated tau species decreased in response to KYNA-1 treatment (pSer202/pThr205 control: 0.94 ± 0.22 KYNA-1: 0.41 ± 0.09, n=7, p=0.044; pThr181 control: 3.35 ± 0.23, KYNA-1: 2.34 ± 0.15, n= 6; p= 0.0035) (Figure 9b,c and d). Our results showed that KYNA-1 in presence of astrocytes can significantly modulate phosphorylation of tau protein. The results also point to important role of astrocytes in abnormal modificantions of tau protein.

## 4 DISCUSSION

The KP is the primary pathway for the metabolism of the essential amino acid tryptophane. During neuroinflammation, tryptophan is catabolized through KP to several neuroactive metabolites, such as KA. In the present study, we explored the effect of synthetic analog of KA (KYNA-1) on the progression of neurofibrillary pathology in a transgenic rat model for tauopathies. Our main findings were as follows: (i) KYNA-1 penetrates BBB better than kynurenic acid, (ii) KYNA-1 reduces the phosphorylation of tau on pThr181 and pSer202/pThr205 epitopes, (iii) chronic administration of KYNA-1 reduces tau phosphorylation and astrogliosis in a transgenic rat model for tauopathies, (iv) KYNA-1 reverse LPS-induced inflammatory changes in glial cell cultures and stimulates anti-inflammatory M2 phenotype in microglia.

BBB represents a bottleneck for the delivery of drugs into the brain. To increase the BBB permeability, small molecules can either be converted into prodrugs or chemically modified [30]. Here we showed that chemical modification of KA (amidation of carboxy group) increased permeability of KA through the *in vitro*BBB model several times. Furthermore, KYNA-1, the N-dimethylaminoethyl amid of KA showed very low toxicity in neuroblastoma cells and primary rat astrocytes. This is in good agreement with studies using the coronal brain slices [22].

In this study, KYNA-1 showed a short half-life of 2.6 h in rats. KYNA-1 was rapidly metabolized into the KA. The concentrations in brain tissue compared to plasma were significantly lower (50 times). This suggests limited BBB permeability in comparison to common CNS active drugs [31]. However, due to a good safety profile, repeated dosing and higher dosage was possible. Moreover, several stable and more BBB permeable analogs were described recently [32]. We also demonstrate increased plasma levels of anthranilic acid and decreased levels of xanthurenic acid after KYNA-1 administration. In the kynurenine pathway, kynureninase regulates the conversion of kynurenine to anthranilic acid [33]. Our data suggests, that when there is an abundance of KA, the KP cascade shifts towards the production of anthranilic acid. Such competition has been mentioned previously [34].

Tauopathies are complex and progressive disorders characterized by brain accumulation of hyperphosphorylated tau protein in intracellular neurofibrillary tangles, neuronal loss, and chronic inflammation. The hyperphosporylated tau from AD brain is most notably stained with AT8 (pS202/pT205), pT217, AT100 (pT212/pS214) and PHF-1 (pS396/pS404) antibodies [35]. Tau phosphorylation on specific epitope before filament formation suggest that hyperphosphorylation is important for conformational changes that trigger tau aggregation and toxicity [36]. Recently, plasma tau has been proposed as an ideal biomarker for AD that overcome the problems with cerebrospinal fluid collection [37].

In the current study, we showed that chronic three month administration of KYNA-1 significantly reduced insoluble hyperphosporylated tau in the brains of transgenic animals. In addition we detected reduced plasma tau levels in KYNA-1 treated animals. No change was observed in the levels of soluble tau, thus we conclude that the reduction is rather indicator of increased degradation of tau by glial cells than due to inhibition of phosporylation. This was further confirmed in multicellular model where only soluble tau is present in NB cells. The co-cultivation of NB with glia was neccessary to induce the KYNA-1 effect.

Neuroinflammation in tauopathies is associated with the activation of microglia and astrocytes along with increased levels of pro-inflammatory mediators such as cytokines and chemokines. Various kynurenines have a biological importance due to their ability to modify neurotransmission and immune response. KA acts as endogenous neuroprotectant and abnormalities in its synthesis have been implicated in the pathogenesis of neurodegenerative disorders. KA is one of the most important metabolite of KP in terms potential therapeutic role [38].The accumulation of reactive astrocytes is one of the hallmarks of tauopathies. They surround plaques and tangles in AD [39]. The characterization of reactive astrocytes is often based on the expression of GFAP. Increased expression of GFAP has been described in tauopathies with predominant tau pathology such as PSP, CBD and PD [40]. Our study provides the evidence that treatment with KYNA-1 significantly reduced the increased GFAP expression in the brain areas affected by tau pathology. To further investigate the mechanism of neuroprotective properties of KYNA-1 analog, we studied the effect of KYNA-1 treatment on experimentaly induced neuroinflammation in cultured astrocytes. Immunoblotting revealed that KYNA-1 decreased expression of GFAP induced by LPS treatment. Another effect was a decrease in toxicity and nitric oxide levels. Notably, the cells retained the non-reactive phenotype.

In the human brain, tau pathology follows distinct pattern. Several studies proved that tau pathology spreads from the localy agregated "seed” through the brain [41]. It has been showed that reactive astrocytes can actively contribute to this process after internalization of pathological tau forms [42]. Hence, we can speculate that the beneficial effect of KYNA-1 on tau pathology in our animal model may be partly attributed to modulation of tau spread between neurons.

Microglia are highly dynamic cells that contribute to healthy function of the CNS. Dysfunctional microglia help to spread pathological tau through phagocytosis and subsequent release of tau-containing exosomes [43]. Accumulating evidence suggests that two major types of activated microglia exists [44]. The M1 phenotype is a classical activation phenotype that is associated with the release of pro-inflammatory cytokines and reactive oxygen species. The M2 activation phenotype is mostly associated with repair processes During the M2 activation microglia release anti-inflammatory cytokines, neurotrophic factors and processes the cell debris [45]. Thus, alteration of the M1/M2 ratio may become an interesting therapeutic intervention in neurodegeneration. In the current study, KYNA-1 induced more M2 anti-inflammatory phenotype that was characterized by reduced expression of CD16 and CD32 and increased expression of CD163. Investigation of the whole signalling cascade involved may provide deeper understanding of anti-inflammatory properties of KYNA-1.

Altogether our results strongly suggest that KYNA-1 has a neuroprotective effect and can reduce pathological tau hyperphosporylation through the induction of anti-inflammatory phenotype in astrocytes and microglia.

## Abbreviations

KP: kynurenine pathway
KA: kynurenic acid
CNS: central nervous system
NMDA: N-methyl-D-aspartate
GABA: γ- aminobutyric acid
LPS: lipopolysaccharide
NO: nitric oxide
AD: Alzheimer’s disease
BBB: blood-brain barrier
KYN: kynurenine

## ACKNOWLEDGEMENTS

This project was supported by grants from the International Centre for Genetic Engineering and Biotechnology (CRP/19/016) and Slovak Research and Development Agency (APVV-18-0302, APVV-16-0531, APVV-18-0259).

## AUTHOR CONTRIBUTIONS

P.M., J.V., A.M and A.K. performed animal studies. P.M. and M.B. performed cell culture experiments and immunocyto-and histochemistry. D.O., G.G., J.P. performed LC/MS experiments. J.H. and P.M. performed biochemical analysis. P.M., D.O. and A.K. prepared the manuscript. All authors have given approval to the final version of the manuscript.

