## Supplementary data for "Analog of Kynurenic Acid Decreases Tau Pathology by Modulating Astrogliosis in Rat Model for Tauopathy"

### (Supplementary Material)

**Table of content**

**Table S1.** Accuracy, precision and recovery of the UHPLC-MS/MS method

**Table S2.** Linearity parameters

**Table S3** – Plasma and brain concentrations of KYNA-1 and KP metabolites after chronic administration to SHR-24 rats

**Figure S1** - Characterization of the multi-component cellular model system.

**Table S1.** Accuracy, precision and recovery of the UHPLC-MS/MS method

RAT PLASMA

| | | Intra-day (n= 5) | | | Inter-day (n= 3) | | |
|---|---|---|---|---|---|---|---|
| **Analyte** | **Spiked concentration** | **Mean concentration** | **RSD (%)** | **Accuracy (%)** | **Mean concentration** | **RSD (%)** | **Accuracy (%)** |
| Anthranilic acid | QC L | 4.90 | 3% | 98% | 5.3 | 0.06 | 107% |
| | QC H | 53.80 | 3% | 108% | 49.7 | 0.11 | 99% |
| Kynurenic acid | QC L | 41.50 | 4% | 83% | 40.8 | 0.08 | 82% |
| | QC H | 423.40 | 3% | 85% | 411.4 | 0.05 | 82% |
| Kynurenine | QC L | 6.50 | 15% | 131% | 5.4 | 0.16 | 108% |
| | QC H | 51.90 | 5% | 104% | 46.8 | 0.13 | 94% |
| KYNA-1 | QC L | 222.40 | 9% | 89% | 214.6 | 0.11 | 86% |
| | QC H | 2315.10 | 4% | 93% | 1946.5 | 0.29 | 78% |
| Xanthurenic acid | QC L | 21.30 | 5% | 85% | 19 | 0.10 | 76% |
| | QC H | 200.10 | 7% | 80% | 189.5 | 0.15 | 76% |

RAT BRAIN TISSUE

| | | Intra-day (n= 5) | | | Inter-day (n= 3) | | |
|---|---|---|---|---|---|---|---|
| **Analyte** | **Spiked concentration** | **Mean concentration** | **RSD (%)** | **Accuracy (%)** | **Mean concentration** | **RSD (%)** | **Accuracy (%)** |
| Anthranilic acid | QC L | 10.00 | 1% | 103% | 10.8 | 0.02 | 103% |
| | QC M | 49.60 | 1% | 111% | 50.9 | 0.03 | 112% |
| | QC H | 99.50 | 1% | 113% | 97.7 | 0.02 | 110% |
| Kynurenic acid | QC L | 11.20 | 2% | 121% | 16.4 | 0.03 | 105% |
| | QC M | 45.90 | 3% | 103% | 55.6 | 0.05 | 111% |
| | QC H | 95.40 | 6% | 108% | 100.8 | 0.03 | 107% |
| Kynurenine | QC L | 56.10 | 2% | 85% | 176.1 | 0.01 | 94% |
| | QC M | 113.70 | 2% | 83% | 236.6 | 0.01 | 88% |
| | QC H | 195.10 | 5% | 88% | 320.3 | 0.04 | 92% |
| KYNA-1 | QC L | 155.30 | 5% | 110% | 151.3 | 0.04 | 107% |
| | QC M | 752.00 | 0% | 107% | 713.4 | 0.01 | 102% |
| | QC H | 1444.80 | 0% | 103% | 1377.9 | 0.01 | 98% |
| Xanthurenic acid | QC L | 41.20 | 2% | 113% | 43.1 | 0.03 | 102% |
| | QC M | 112.90 | 4% | 105% | 116.1 | 0.02 | 104% |
| | QC H | 207.00 | 5% | 106% | 214.8 | 0.03 | 108% |

**Table S2.** Linearity parameters

RAT BRAIN TISSUE

| Analyte | Retention time | $R^2$ | Regression equation | Linear range | LOD | LOQ |
|---|---|---|---|---|---|---|
| Anthranilic acid | 2.3 | 0.9997 | y = 0.0896096x + 0.00060822 | 0.1 - 100 | 0.27 | 0.9 |
| Kynurenic acid | 1.9 | 0.9983 | y = 0.0457023x + 0.00091791 | 0.1 - 100 | 0.03 | 0.1 |
| Kynurenine | 1.6 | 0.9984 | y = 0.0358862x + 0.00354954 | 1 - 200 | 0.02 | 0.1 |
| KYNA-1 | 1.7 | 0.9989 | y = 0.0046900x + 0.0113103 | 8 - 1600 | 0.002 | 0.005 |
| Xanthurenic acid | 1.8 | 0.9982 | y = 0.0149696x + 0.00073985 | 1 - 200 | 0.87 | 0.1 |

RAT PLASMA

| Analyte | Retention time | $R^2$ | Regression equation | Linear range | LOD | LOQ |
|---|---|---|---|---|---|---|
| Anthranilic acid | 2.3 | 0.9989 | y = 0.0181669x + 0.0032427 | 0.5 - 100 | 0.03 | 0.12 |
| Kynurenic acid | 1.9 | 0.9997 | y = 0.0240833x + 0.0003379 | 0.5 - 100 | 0.01 | 0.03 |
| Kynurenine | 1.6 | 0.9990 | y = 0.0496237x + 0.0007186 | 5 - 1000 | 0.04 | 0.14 |
| KYNA-1 | 1.7 | 0.9990 | y = 0.0059996x + 0.0207487 | 25 - 5000 | 0.02 | 0.08 |
| Xanthurenic acid | 1.8 | 0.9992 | y = 0.0232125x + 0.0001841 | 2.5 - 500 | 0.01 | 0.03 |

**Table S3** – Plasma concentrations of KYNA-1 and KP metabolites after chronic administration to SHR-24 rats

| | | KYNA-1 chronic administration | Placebo group | |
|---|---|---|---|---|
| | **Compound** | Mean ± SD (ng/g; ng/ml) | Mean ± SD (ng/g; ng/ml) | |
| **Brainstem** | **Anthranilic acid** | 3.17 ± 0.39 | 3.56 ± 0.63 | ns |
| | **Kynurenic acid** | 0.64 ± 0.28 | 0.96 ± 0.55 | ns |
| | **Xanthurenic acid** | 28.08 ± 3 | 29.59 ± 2.42 | ns |
| | **Kynurenine** | 31.8 ± 6.55 | 36.32 ± 8.33 | ns |
| | **KYNA-1** | 97.09 ± 144.9 | - | |
| **Plasma** | **Anthranilic acid** | 7.6 ± 0.9 | 9.2 ± 2.5 | ns |
| | **Kynurenic acid** | 14.5 ± 5.9 | 21.6 ± 4.8 | p=0.011 |
| | **Xanthurenic acid** | 31 ± 8.6 | 40.6 ± 7.6 | p=0.02 |
| | **Kynurenine** | 210.3 ± 67.9 | 268 ± 62.9 | ns |
| | **KYNA-1** | 912.9 ± 1208 | - | |

ns – p>0.05

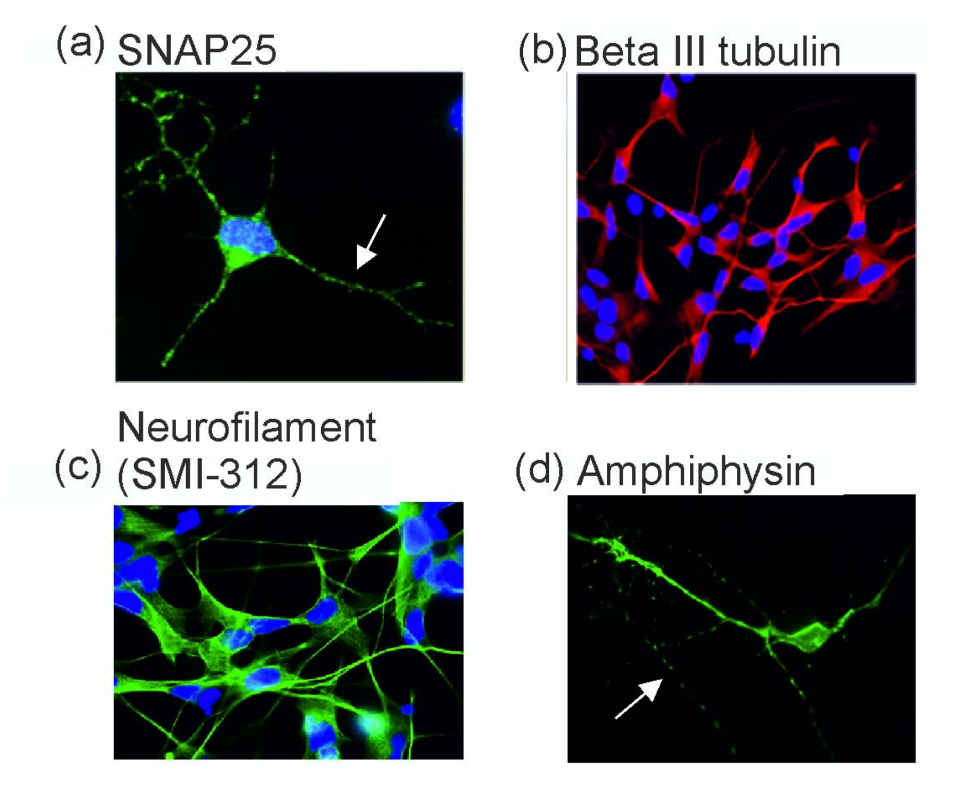


**Figure S1 - Characterization of the multi-component cellular model system.** The results from immunocytochemical staining showed the re-distribution of neuron-specific and synaptic proteins from the cellular body into cell projections (white arrow). Differentiated neuroblastoma cells expressed neuronal-specific markers after co-cultivation with primary rat astrocytes: SNAP25 (green, a), Beta III tubulin (red, b), Neurofilament SMI-312 (green, c), amphiphysin (green, d). DAPI was used as nuclei staining (blue). Scale bar: 20 μm.
